# Chromatin Accessibility Shapes Developmental-Specific Lineage Plasticity in Hematopoiesis

**DOI:** 10.64898/2025.12.02.691283

**Authors:** Sara Palo, Keiki Nagaharu, Mikael Sommarin, Rasmus Olofzon, Virginia Turati, Shamit Soneji, Göran Karlsson, Charlotta Böiers

## Abstract

Cell differentiation is governed by dynamic changes in chromatin accessibility, and its dysregulation underlies multiple disease states. Prior to birth, development of the hematopoietic system constitutes a period of broad differentiation potential, with certain immune cells arising exclusively during ontogeny. While age is known to affect lineage bias, the underlying molecular differences driving lineage preference in fetal and adult human hematopoietic stem and progenitor cells (HSPCs) remain unclear. Through single-cell cultures of hematopoietic stem cells (HSCs), we observed that fetal cells frequently generate mixed-lineage colonies, whereas adult HSCs are largely restricted to myeloid output. To investigate how these lineage preferences were encoded at the chromatin level, we performed single-cell ATAC-sequencing on first-trimester HSPCs. While adult HSCs showed enrichment of lineage-specific transcription factor motifs, fetal cells lacked such enrichment, consistent with their broader differentiation potential. We additionally uncovered a developmental-specific plasticity in fetal lymphoid progenitors, manifested as a hybrid lympho-myeloid chromatin program not present in adult progenitors. Additionally, the motif for PAX5, master regulator of B cell development, showed markedly reduced accessibility in fetal cells, indicating a more plastic and less committed lymphoid state. This enhanced embryonic lineage plasticity may underlie the prenatal susceptibility to mutational drivers of acute lymphoblastic leukemia.

## Introduction

During ontogeny, stem and progenitor cells possess broad differentiation potential, with certain immune cell types arising exclusively during fetal development^1^, a flexibility likely necessary for the establishment of our complex immune system. Transcription factors (TFs) are key regulators of the differentiation process: by binding to lineage-associated gene promoters and enhancers, they are essential to the formation of cell identity. Complementary to TF binding, changes in chromatin accessibility reshape the epigenomic landscape and have been suggested to prime cells for subsequent lineage commitment by enabling or restricting transcription of specific regions of the genome^2, 3^. Consequently, differences in TF motif accessibility have been connected to lineage affiliation^4, 5^.

Within the hematopoietic system, self-renewing, multipotent hematopoietic stem cells (HSCs) generate increasingly lineage-restricted progenitors that proceed along differentiation trajectories toward mature cell types. The advent of single-cell technologies has uncovered heterogeneity within these immunophenotypically discrete cell states, revealing a continuum of gradually more restricted progenitor cells^6–8^. In parallel, functional studies have demonstrated that HSCs become increasingly myeloid and platelet-biased with aging^9–12^. Ultimately, this age-dependent shift in lineage bias mirrors the occurrence of acute leukemias, where lymphoid leukemias predominate in children and myeloid leukemias in aged individuals^13^.

Despite age being a well-known factor affecting lineage bias, most studies of fetal hematopoietic lineage commitment have focused solely on ontogeny, with few comparisons to the adult^5, 14–18^. During ontogeny the blood system develops over several phases, or waves, and across multiple niches^19, 20^. In a reversal of the typical hierarchy, definitive HSCs emerge last, in the aorta-gonad-mesonephros (AGM) region starting at around 30 days post conception (Carnegie stage (CS) 14) and then migrate to the fetal liver (FL), the main niche for expansion and maturation of hematopoietic cells during early fetal life. In the bone marrow (BM), which will maintain hematopoiesis throughout adulthood, hematopoiesis becomes evident at the beginning of the second trimester^19, 20^. The heterogenous composition of the blood system necessitates the use of single-cell assays. While fetal hematopoietic stem and progenitor cells (HSPCs) have been shown to differ from adult counterparts in composition, cell cycle status and transcriptional programs^15, 16, 21^, lineage priming is challenging to assess at the single-cell transcriptional level. Transcription factors (TFs) are central regulators of differentiation and crucial for establishing cell identity, but due to their often-low expression, TFs often remain undetected in single-cell RNA sequencing. Chromatin-level studies can assess chromatin accessibility at TF binding motifs, but the few published to date do not include direct comparisons with adult HSPCs^5, 22, 23^. Consequently, it remains unclear whether lineage priming and differentiation pathways during ontogeny diverge from those observed in adulthood.

To address this knowledge gap, we performed parallel single-HSC cultures to assess differences in lineage preference across development and profiled the chromatin landscape of first trimester human fetal liver HSPCs at single-cell resolution. By directly comparing to adult counterparts, our analyses uncover developmental stage-dependent, chromatin-encoded differences in lineage bias and plasticity of the hematopoietic system.

## Results

### Hematopoietic stem cell lineage bias is developmental stage-dependent

We first investigated the effect of developmental origin on HSC lineage preference *in vitro*. To capture different stages of development we selected samples representing first trimester FL, the main site of hematopoietic expansion and maturation during early ontogeny^19^; neonatal blood (cord blood (CB)) and adult BM. To accurately assess lineage preference, cultures were performed at the single cell level in conditions designed to promote multi-lineage differentiation. Single HSCs (defined as lineage negative (LIN^-^)CD45^+^CD34^+^CD38^-^CD90^+^CD45RA^-^)^24^ were index sorted and cultured for three weeks, after which output was assessed by flow cytometry (**Figure 1A, Figure S1A-B**). The colonies generated from adult HSCs were mainly myeloid, with minor contributions of other lineage types. Fetal-derived colonies frequently exhibited erythroid output, as previously reported^14^. However, in contrast to an earlier report that observed no B cell output until the second trimester^14^, we found robust B-lineage potential in first trimester HSCs and nearly one-third of the colonies displayed lympho-myeloid lineage differentiation. Neonatal HSCs exhibited an intermediate phenotype, with a higher proportion of single myeloid colonies than FL, while retaining the capacity to generate lympho-myeloid lineage colonies (**Figure 1B-E, Figure S1C-E**). Overall, fetal HSCs exhibited mixed lineage output, with 43% of the colonies displaying bi-or oligo-lineage output, whereas a majority (∼70%) of the colonies derived from adult HSCs were restricted to a single lineage **(Figure 1F, Figure S1F).**

**Figure 1.**
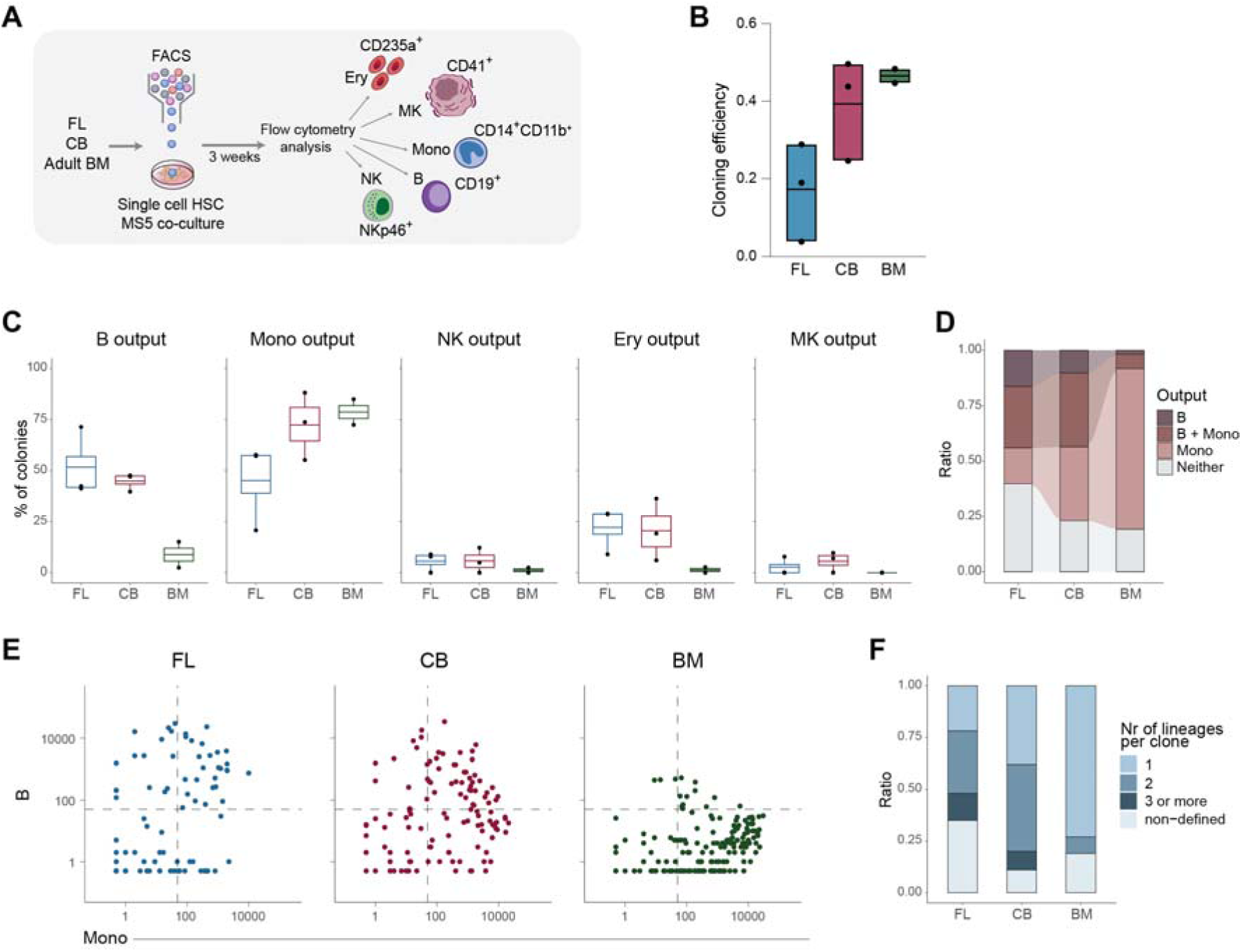
Hematopoietic stem cell lineage bias is developmental stage-dependent. **(A)** Schematic illustration of *in vitro* culture experiments and markers used to define lineage identity. **(B)** Cloning efficiency as percentage of sorted HSCs. Each dot represents one experiment, and the line mean value (n_FL_ = 3 (∼7-9pcw), n_CB_ = 3, n_BM_ = 2 (∼20-30 years old). A fourth FL sample produced no colonies and was excluded). **(C)** Lineage output presented as the percentage of total number of colonies, displayed per donor for each lineage. The line shows the mean value. **(D)** Proportions of colonies with output of B but no Mono, B and Mono, Mono but no B, or neither of these lineages, for all replicates combined. **(E)** B and Mono lineage output displayed per colony. Dashed lines indicate the 50-cell cutoff for lineage positivity. Zero counts were assigned a value of 0.5 to account for the logarithmic scale of the axes. **(F)** Proportions of colonies with output of one, two or ≥3 defined lineages, or no defined lineage, for all replicates combined.

These *in vitro* culture results reveal a clear developmental regulation of HSC lineage preference, with first trimester HSCs exhibiting enhanced B lineage output and broader lineage flexibility relative to adult HSCs.

### The fetal HSPC chromatin landscape reveals closely connected cell states

To investigate if the developmental differences in lineage preference observed *in vitro* were encoded at the chromatin-level, we performed single-cell assay for transposase-accessible chromatin using sequencing (ATAC-seq). To ensure comprehensive coverage of the primitive compartment, LIN^-^CD45^+^CD34^+^ HSPCs were enriched from two CS22 (∼8 post-conceptional weeks (pcw)) donors (**Figure 2A, Figure S2A**, **Table S1)**. The cells (>13,000) were integrated using Harmony^25^, and the subsequent uniform Manifold Approximation and Projection (UMAP) visualization resulted in 14 clusters (**Figure 2B, Figure S2B-C**). Cell states were assigned based on accessibility of transcription factor (TF) motifs^26^, with HOXB motifs in cluster #0 indicating an HSC identity^4^, while EBF1 in cluster #4, CEBPA in cluster #8 and GATA1 in cluster #13, denote B-lymphoid (Ly), myeloid (My) and megakaryocyte-erythroid (MkE) cell states, respectively^27^ (**Figure 2C-D**).

**Figure 2.**
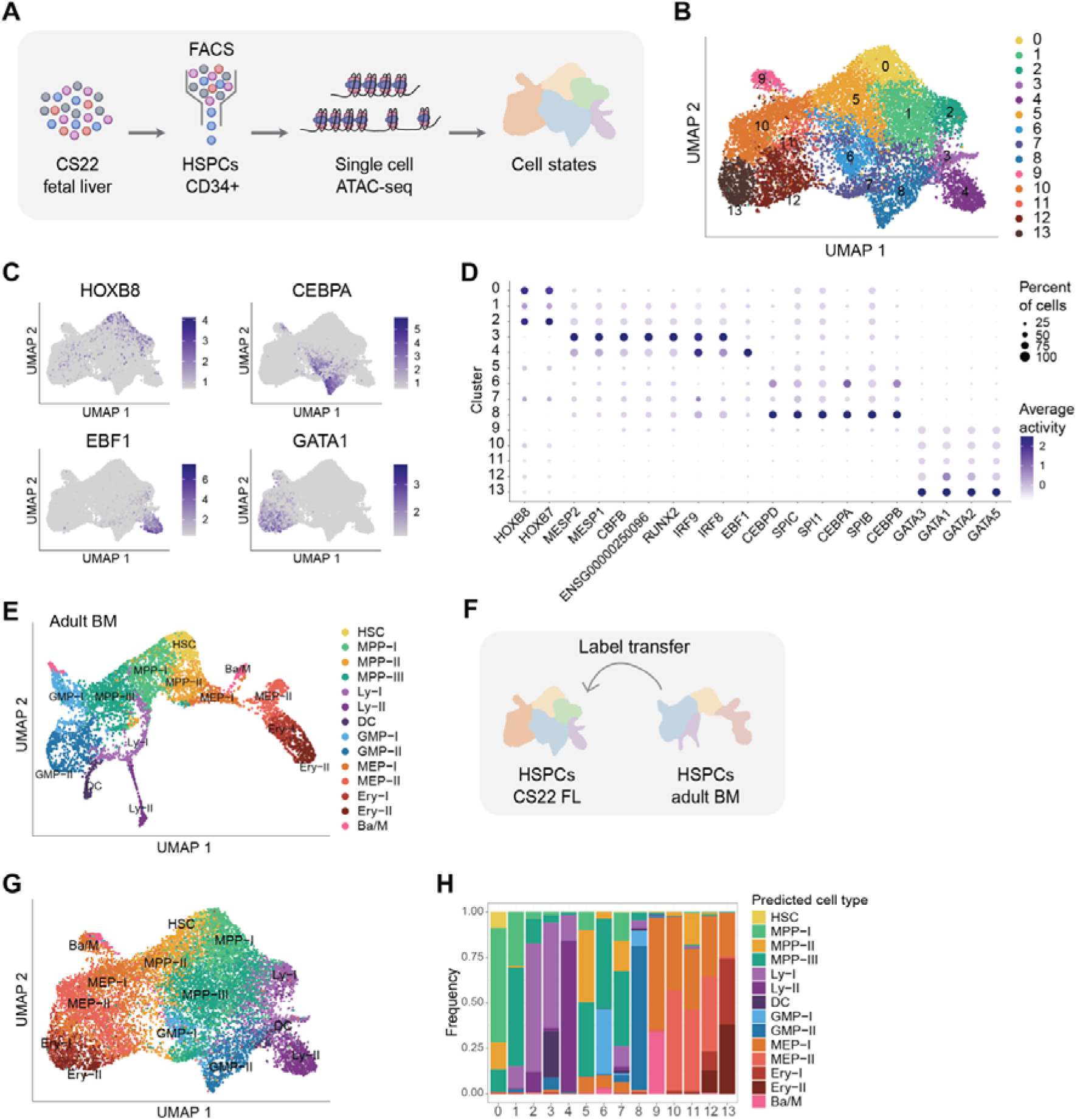
The fetal HSPC chromatin landscape reveals closely connected cell states. **(A)** Schematic illustration of the experimental workflow. **(B)** UMAP of CS22 FL HSPCs (n = 2 donors, total 13,953 cells). (**C**) Motif activity for different lineage-associated transcription factors. The top 50% of cells are colored by activity. (**D**) Dot plot of the motifs with highest differential activity between clusters (mean z-score difference > 1.5; p-adj < 0.05). (**E**) UMAP of adult BM HSPCs from^28^. **(F)** Schematic illustration of the label transfer. **(G)** The FL UMAP from (B) annotated with predicted cell identities from the label transfer. **(H)** Distribution of predicted cell identities from the label transfer per FL Seurat cluster.

To enable direct comparison of chromatin accessibility between fetal and adult HSPCs, we utilized our previously published dataset of LIN^-^CD34^+^ adult BM HSPCs^28^. The adult BM UMAP separated into distinct lineage-associated clusters, displaying two prominent branches: one comprising lympho-myeloid clusters and the other consisting of megakaryocytic-erythroid clusters. In contrast, the FL HSPCs formed a more continuous landscape, with hematopoietic lineages appearing closely connected, suggesting a less compartmentalized state during development (**Figure 2B, 2E, Figure S2D-E**). We next projected the FL cells onto the adult BM UMAP and annotated them based on their predicted adult cell type, which was in line with the TF motif activity (**Figure 2F-H, Figure S2F**). Compared to the adult BM, a smaller proportion of FL cells were annotated as HSCs, and a higher fraction as lymphoid, in agreement with what has previously been shown at the transcriptional level^15, 21^ (**Figure S2G**).

Thus, while first-trimester HSPCs separated into cell states based on chromatin accessibility, these cell states appeared more interconnected and less distinct than their adult counterparts, potentially reflecting a higher degree of lineage plasticity.

### Limited lineage-specific chromatin priming in first trimester HSCs

Utilizing the aforementioned datasets, we aimed to resolve lineage priming in HSCs at the chromatin-level. To cover a larger span of ontogeny, we incorporated a publicly available dataset of second trimester (18-21 pcw) FL and fetal BM HSPCs^5^ (**Figure 3A**). We annotated the cells based on projection to the adult BM data as described for the first trimester dataset, and then calculated differentially accessible peaks between each of the three fetal tissues and adult BM. In general, there were more up-than downregulated peaks prenatally compared to adult, independent of fetal stage (**Figure 3B, Figure S3A-B, Table S2**). As TFs drive lineage commitment and changes in TF motif activity are early indicators of priming^4, 5^, we searched for enriched TF motifs within these regions. During prenatal development, only a few TF motifs were found to be enriched. In the first trimester, these included well-known regulators of HSC self-renewal, such as HOXB7 and HOXB8. By the second trimester, SPI-B, a member of the SPI family of transcription factors, was observed. SPI-B is closely related to SPI-1 (PU.1) and acts mainly in B cell differentiation, while SPI-1 functions more broadly in lympho-myeloid lineage specification^29, 30^. In contrast, myeloid and MkE-associated motifs, such as JUN, FOSL and NF-E2 were enriched in adult BM^31, 32^ (**Figure 3C-D, Figure S3C, Table S3**). For many of these motifs we also detected, based on footprinting analysis, a difference in TF occupancy between fetal and adult HSCs, with e.g. JUN and NF-E2 signals being higher in the adult. SPI-B, which was enriched in the peaks upregulated in second trimester FL HSCs, also exhibited the strongest footprint in second trimester cells, independent of niche (**Figure 3E, Figure S3D**).

**Figure 3.**
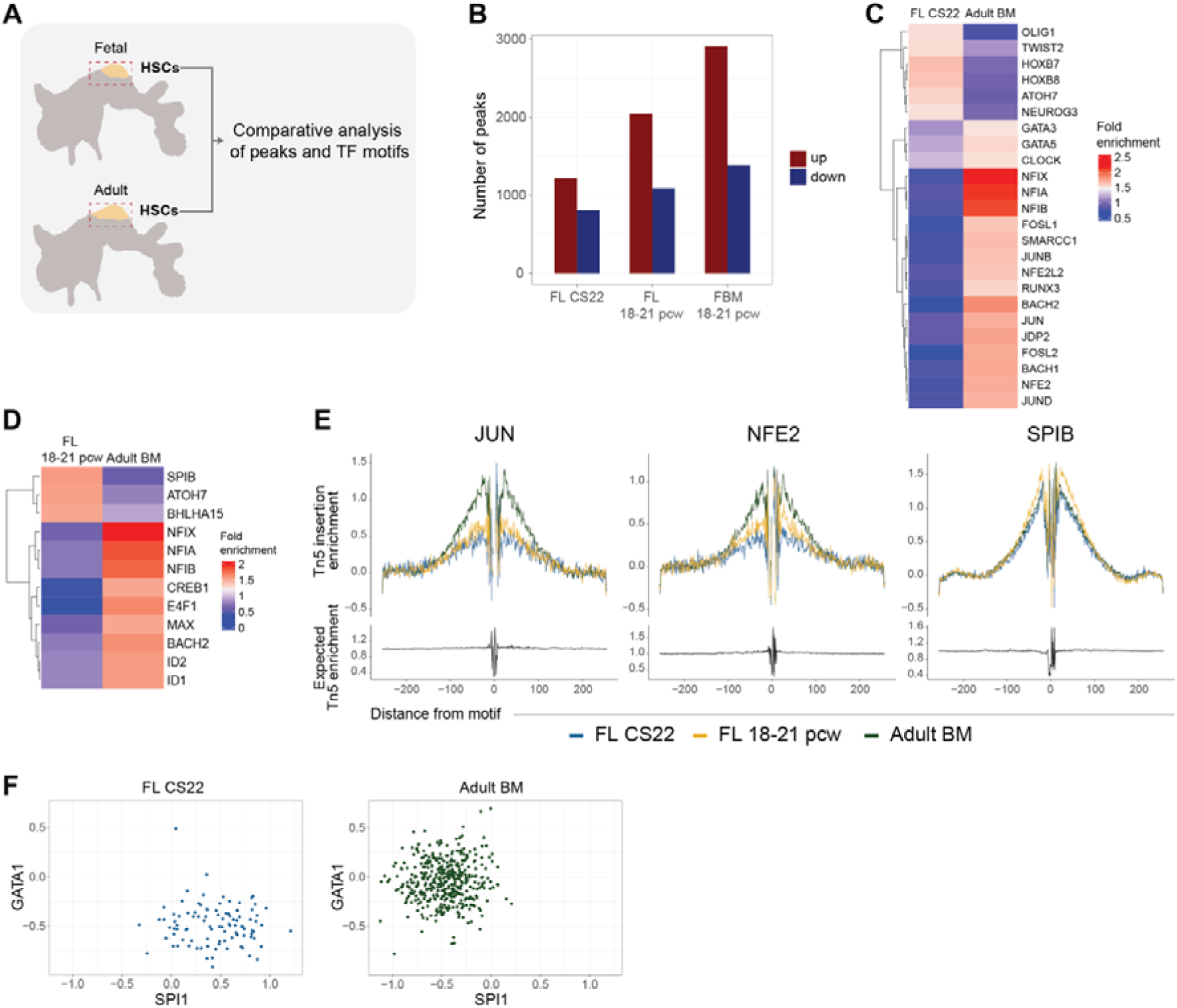
Limited lineage-specific chromatin priming in first trimester HSCs. **(A)** Schematic illustration of the analysis. **(B)** Number of differentially accessible peaks between HSCs from different fetal tissues and adult BM HSCs (log2 fold change > 2; p-adj < 0.05). **(C)** Heatmap of TF motifs enriched (fold change > 1.5, p-adj < 0.05) in peaks differentially accessible between CS22 FL and adult BM HSCs. **(D)** As (C) but for second trimester FL^5^ and adult BM HSCs. **(E)** Footprints of selected TFs from (C-D), for FL and adult BM HSCs. **(F)** Motif activity for GATA1 and SPI1 in CS22 FL *(left*) and adult BM (*right*) HSCs. The activity values were scaled across the entire datasets prior to subsetting for HSCs.

We next focused on GATA1 and SPI-1 (PU.1), two TFs known to play key roles in hematopoietic cell fate decisions. GATA1 act as a master regulator of erythroid differentiation, and SPI-1 orchestrates myeloid and lymphoid cell fates as noted above^29, 33, 34^. Our analysis found clear developmental stage-based differences in relative accessibility of their motifs, with SPI-1 relatively more accessible in the first trimester FL HSCs and GATA1 in adult (**Figure 3F**).

Taken together, our results suggest limited lineage priming of the HSC state during the first trimester, allowing for a diverse output of mature blood cells, whereas adult HSCs have myeloid lineage priming encoded at the chromatin level, in agreement with the myeloid lineage bias observed *in vitro* (**Figure 1C-F**).

### The fetal lymphoid cell state exhibits a lympho-myeloid chromatin signature

The developmental differences in lineage priming and lineage output at the HSC level prompted us to explore changes in chromatin accessibility as cells transition into more differentiated cell states, focusing our analysis on the lymphoid and myeloid clusters. We first investigated how TF motif accessibility changes between HSCs and the most mature progenitor states in our fetal and adult datasets (**Figure 4A**). The fetal lymphoid cluster (Ly-II) was heavily biased towards a gain of accessible motifs, while very few motifs were lost. Furthermore, we found that these gained TF motifs had a lympho-myeloid lineage affiliation, whereas in the adult cells, the motifs with increased activity had a more distinct lymphoid profile. A similar pattern was observed in the more immature Ly-I cluster. In contrast, the TF motifs gained in the myeloid clusters (GMP-I and GMP-II) were associated with the myeloid lineage, irrespective of developmental stage (**Figure 4B, Figure S4A-B, Table S4**).

**Figure 4.**
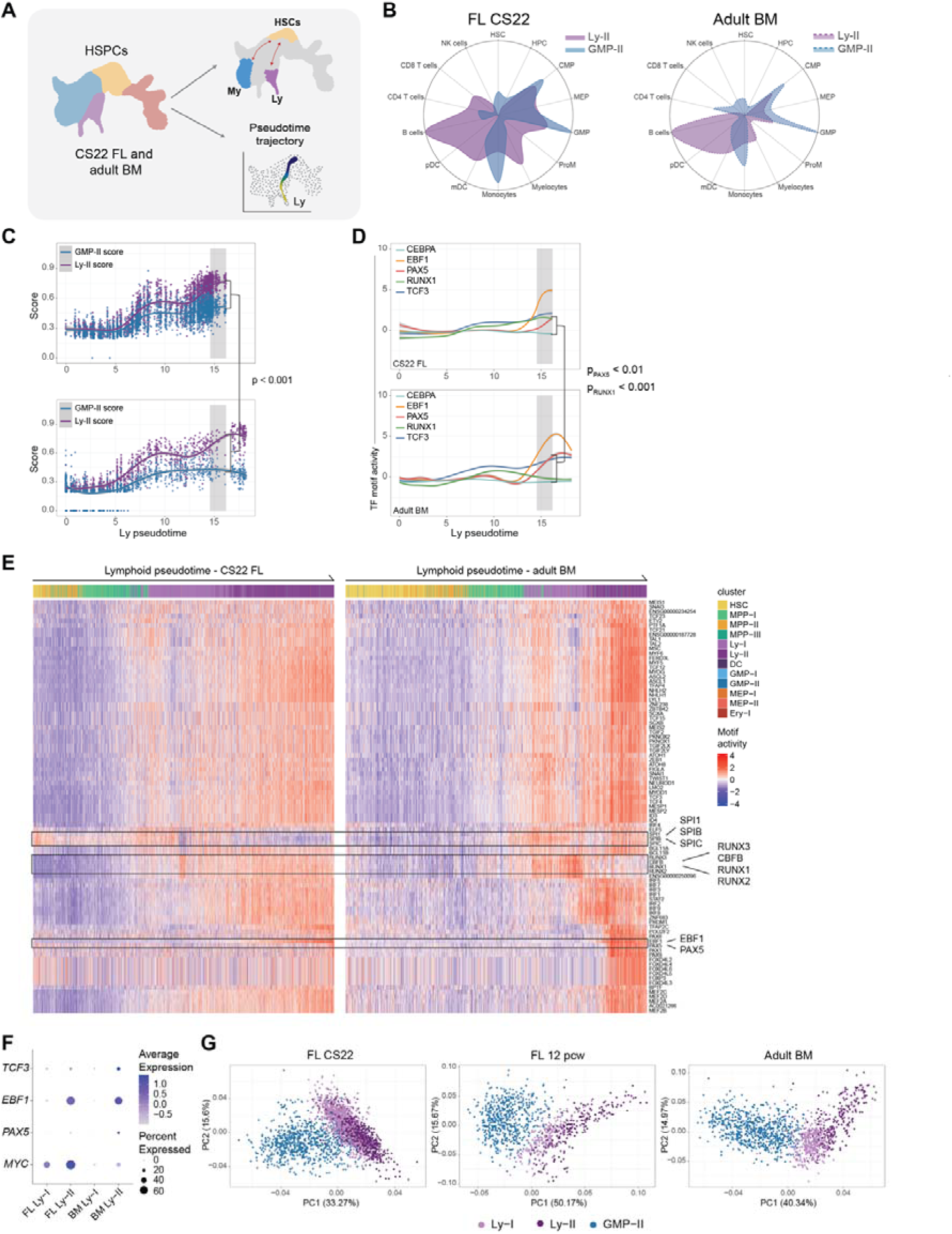
Lympho-myeloid chromatin plasticity and reduced B lineage commitment in early fetal lymphoid progenitors. **(A)** Schematic illustration of the analysis. **(B)** Lineage affiliation of TF motifs with increased activity compared to the HSC cluster for the Ly-II and GMP-II clusters for CS22 FL (*left*) and adult BM (*right*). **(C)** Scores for TF motifs upregulated in adult BM Ly-II or GMP-II displayed along the lymphoid trajectory for the FL (*upper*) and adult BM cells (*lower*). (p < 0.001, *see method section and Figure S4E*). **(D)** Activity of B cell commitment-associated TF motifs along the Ly trajectory for FL (*upper*) and BM (*lower*). CEBPA is shown as a reference (PAX5 p_adj_ <0.01, and RUNX1 p_adj_ < 0.001, *see method section and Figure S4F*). **(E)** Heatmap of motif activity along the lymphoid trajectory for motifs upregulated in Ly-I or Ly-II in FL or adult BM, compared to HSCs in FL (*left*) or adult BM (*right*). **(F)** mRNA expression of selected genes in first trimester FL^21^ and adult BM^28^ lymphoid progenitors. **(G)** PC analysis based on the top 150 variable TF motifs for Ly-I, Ly-II and GMP-II cells for CS22 FL (*left*), 12 pcw FL^22^ (*middle*) and adult BM^28^ (*right*).

To obtain a better understanding of the developmental differences in the lymphoid pathway, we looked more closely at the trajectory from the HSC cell state to the most mature lymphoid cluster and calculated scores for the lymphoid and myeloid upregulated motifs. While the adult trajectory displayed a clear separation between the myeloid and lymphoid signatures, these remained more closely connected throughout the fetal trajectory (**Figure 4C, Figure S4C-E, Table S5**), implying a less committed and more plastic lymphoid cell state during early development.

### Reduced chromatin-level commitment to the B lineage in fetal lymphoid progenitors

Given the reduced lineage segregation observed in the fetal trajectory, we next asked whether this developmental plasticity is also reflected in the chromatin programs that govern B cell commitment. TCF3 (E2A) precedes expression of EBF1, which in turn controls PAX5^35^, a master regulator of B cell commitment. As expected, the motifs for TCF3 and EBF1 were upregulated compared to HSCs in both fetal and adult cells. In contrast, PAX5 was significantly increased in adult BM cells, but not in fetal lymphoid cells. This implies that FL lymphoid cells are less committed to the B lineage at the chromatin level (**Figure 4D-E, Figure S4F, Table S6**).

Looking further into the FL-specific downregulated TF motifs, we found that members of the SPI family (SPI-1, SPI-B, and SPI-C), had reduced activity in FL lymphoid progenitors, whereas in adult cells they exhibited increased activity compared to HSCs (**Figure 4E, Figure S4G)**. This contrasting pattern suggests a developmental regulation of the SPI family, which is known to play a central role in lympho-myeloid lineage specification. Among FL-specific gained motifs we identified RUNX1, RUNX3 and the RUNX1 co-factor CBFB (**Figure 4D-E, Figure S4F-G**). RUNX1 has been shown to be essential to the early stages of B cell differentiation^36^. The fact that RUNX1 remained accessible in FL lymphoid cells, implies a delay in B cell commitment compared to adult counterparts, in line with the lower PAX5 TF motif activity (**Figure 4D-E**). In addition, EBF1 has been shown to directly upregulate the proto-oncogene MYC, while PAX5 can counteract this effect^37^, a mechanism particularly relevant in the context of leukemogenesis. Single cell RNA-seq data from comparable developmental stages^21^ also revealed elevated MYC expression in fetal lymphoid progenitors (**Figure 4F**), supporting the notion of a less restricted, more proliferative state during fetal B cell development.

Finally, we asked whether the B lineage plasticity observed in the fetal liver was specific to the first trimester or reflected a broader prenatal feature. To address this, we leveraged publicly available data from early second trimester (12 pcw) FL^22^ (**Figure S4H-I,** *see method for details*). Performing principal component analysis (PCA) on the top variable motifs in each dataset, we found that the first trimester Ly-II cells clustered closely together with the less mature Ly-I cells, suggesting a more plastic and less restricted lymphoid state. In striking contrast, in second trimester FL and adult BM the Ly-I and Ly-II populations were clearly separated (**Figure 4G**), indicating a more committed lymphoid progenitor at later developmental stages.

Altogether, these findings show that first-trimester lymphoid progenitors possess a uniquely flexible chromatin state, characterized by reduced B cell priming.

## Discussion

Although transcriptional differences between fetal and adult hematopoiesis have been broadly characterized^15, 16, 21^ the difficulty of reliably detecting transcription factors using single-cell RNA sequencing underscores the importance of chromatin profiling to resolve lineage priming. By performing parallel cultures of HSCs from different developmental stages, we uncovered striking differences in lineage preference. Fetal HSCs exhibited mixed lineage output, whereas adult HSCs predominantly produced single-lineage myeloid colonies. This functional difference was mirrored at the chromatin level: fetal HSCs displayed a broadly permissive chromatin landscape with minimal lineage priming, while adult HSCs showed enrichment of myeloid-associated TF motifs. Although increased Mk potential has been associated with aging in both mice and humans^9, 11, 12^, and some Mk-related motifs such as NF-E2^31^ were enriched, this did not translate into functional Mk output. Given that our adult donors were young it is plausible that chromatin-level Mk priming emerges well before functional bias becomes evident. A further point of interest is the prominent B cell output from early fetal HSCs, contrasting a prior study where B cell output was not observed until in the second trimester^14^. To our knowledge, no prior study has directly compared single human fetal and adult HSCs in multilineage-promoting culture conditions that facilitate B lineage differentiation, and our results uncover a substantial developmental difference in B-Mono output.

The diverse lineage output observed in our first-trimester cultures aligns with the broadly permissive chromatin landscape of early fetal HSCs. Among the few motifs enriched in first-trimester FL HSCs, some were associated with self-renewal regulators. HOX cluster genes are well-known targets of KMT2A fusion-driven leukemias, which dominate among infant patients^38^. The greater accessibility of these TF motifs in FL HSC compared to adult may indicate an increased susceptibility to fusion-mediated transformation during early development.

We also found lineage plasticity in the fetal lymphoid trajectory, seen as lympho-myeloid priming at the chromatin level. We did, however, not observe a lymphoid signature in the myeloid clusters; thus, the flexibility seems specific to the lymphoid progenitors. Of note, the fetal lymphoid progenitor state mainly exhibited a gain of accessible TF motifs, with only a few lost compared to HSCs. This overall increase in motif accessibility may contribute to the lineage plasticity observed. Considering that second trimester progenitor cells have previously been shown to be more oligopotent within erythro-myeloid lineages than adult counterparts^18^, this implies an overall higher lineage plasticity during fetal hematopoiesis.

Pediatric leukemia is known to be driven by different genetic aberrations than adult, and several of these initiating mutations have been shown to arise *in utero*^13, 39^. This suggests that the fetal hematopoietic context allows for certain leukemic driver mutations to occur and/or persist. The lineage plasticity observed in the FL HSPCs could be of importance for the susceptibility to certain leukemia driver mutations that occur during fetal life. One may speculate that a flexible, less restricted chromatin landscape of the initiating cell provides access to target genes in a way that would not be possible in the adult setting. Moreover, the reduced PAX5 motif accessibility we observed along the FL lymphoid trajectory aligns with delayed B-lineage restriction; PAX5 is a critical gatekeeper of B-cell identity^40^. PAX5 haploinsufficiency is a well-recognized secondary mutation in B-ALL^41^, and germline PAX5 variants have been linked to inherited susceptibility to the disease^42^. A less lineage-restricted fetal lymphoid progenitor cell may allow for greater plasticity upon oncogenic insult, enabling malignant clones to access both lymphoid and myeloid programs. Of note, lympho-myeloid lineage switch is a rare, but well-known phenomenon in pediatric B-ALL^43^. While the time window for the occurrence of leukemia-initiating mutations during ontogeny remains unknown, our data suggest that lineage plasticity is primarily a feature of the first trimester. Importantly, early fetal liver hematopoietic cells comprise a heterogeneous mix of transient and emerging definitive populations^19^, and therefore, the observed plasticity cannot be attributed to either transient or definitive hematopoiesis.

Overall, our findings provide new insights into the chromatin architecture of early primitive hematopoietic cells, revealing an ontogeny-specific plasticity. This flexibility, encoded at the epigenetic level, likely underpins the unique developmental potential of fetal hematopoiesis and may contribute to its susceptibility to leukemogenic driver events.

## Methods

### Human samples

Human FLs were donated from elective terminations of pregnancy with informed consent and approval of the Regional Ethics Review Board (Lund)/the Swedish Ethical Review Authority and the Swedish National Board of Health and Welfare. Embryos were staged according to Carnegie staging or pcw. CB and BM aspirates (∼ 20-30 years old) were collected after informed consent and with approval of the Regional Ethics Review Board (Lund)/the Swedish Ethical Review Authority.

FL tissues were dissociated mechanically and frozen in StemCellBanker (Amsbio) or fetal bovine serum (FBS) with 10% dimethyl sulfoxide (DMSO, Sigma). CB and BM samples were diluted (1:1 with PBS (GE lifescience) with the addition of 2% fetal bovine serum (GE lifescience) and 2 mM EDTA (Ambion)) and transferred to lymphoprep tubes (Alare Technologies/Serumwerk/Stemcell technologies), and the mononuclear layer extracted according to manufactureŕs instruction. The mononuclear cells (MNCs) were washed 1:1 with Iscove’s modified Dulbecco’s medium (IMDM, Thermofisher) and were either frozen down or processed further for CD34 enrichment. CD34 enrichment was preformed using CD34 enrichment kit (Miltenyi Biotec) according to manufacturer’s instructions. Briefly, cells were incubated with Fc-block and CD34 enrichment beads for 20 min, after which positive magnetic selection was performed using columns. MNCs or CD34^+^ cells were frozen in FBS with 10% DMSO and stored at −150°C until use.

### Single cell ATAC sequencing preparation

On the day of experiment, FL samples (n = 2, CS22) were thawed and stained with Fc receptor blocking reagent and the following antibodies: CD45-Alexa700, CD34-FITC, CD38-APC and lineage markers (LIN: CD19-BV605, CD3-PE-Cyanine5, CD2-PE, CD16-BV421, CD14-PE-Cyanine7, CD235a-PE-Cyanine5)(see also **Table S7**). LIN^-^CD45^+^CD34^+^ cells were sorted for single cell ATAC-seq using a BD FACSAria IIu. Single cell ATAC-seq was performed using the 10X Genomics single cell ATAC-seq platform according to the manufacturer’s instructions. After library preparation the libraries were sequenced on a NOVAseq (Illumina).

### In vitro cultures

Cryopreserved FL (n = 3, ∼ 7-9 pcw. A fourth FL sample produced no colonies and was excluded.), CB (n = 3) and adult BM (n = 2, ∼ 20-30 years old) samples were thawed, washed, and stained with Fc receptor blocking reagent and the following antibodies: CD45-Alexa700, CD34-APC, CD38-PE-Cyanine7, CD90-BV786, CD45RA-FITC and lineage markers (LIN: CD19-BV605, CD3-PE-Cyanine5, CD14-PE-Cyanine5, CD56-PE-Cyanine5, CD235a-PE-Cyanine5) (see also **Table S7**). 7-AAD was used to assess viability.

The day before sorting, gelatin-coated 96-well plates were seeded with 2000 MS-5 stromal cells per well. Prior to sorting, the culture medium was replaced by 50 μL Myelocult H5100 (Stemcell Technologies) supplemented with 100 ng/mL Stem cell factor (SCF) (PeproTech), 50 ng/mL Fms-like tyrosine kinase 3 ligand (FLT3L) (PeproTech), 50 ng/mL Thrombopoietin (TPO) (PeproTech), 3 U/mL Erythropoietin (EPO) (Retacrit), 20 ng/ml interleukin-6 (IL-6) (PeproTech), 10ng/ml IL-2 (PeproTech), 20 ng/mL IL-7 (PeproTech), and 100 U/mL penicillin-streptomycin (Gibco). Single hematopoietic stem cells (HSCs; defined as Lin^-^ CD45^+^CD34^+^CD38^-^CD45RA^-^CD90^+^) were index sorted into wells with Myelocult using a FACS Aria III (BD). 50 μL medium with 2x cytokine concentration was added on day 3 and 100 μL on day 10.

### Readout and analysis of *in vitro* cultures

All wells were scored under the microscope for colonies and visible colonies were harvested on day 21. The cells were Fc-receptor-blocked and stained for ∼40 min at 4 °C with the following antibodies: CD19-BUV395, CD71-BUV496, CD335-BV421, CD235a-FITC, CD41-PE, CD11b-APC, CD45-Alexa700/PE-Cyanine7, CD14-PE-Cyanine7/Alexa700. 7-AAD was used as viability dye and flow cytometry was performed on a Fortessa X20 (BD). Gating was performed in Flowjo (v10.10). Viable, single cells were gated on size and assigned to a lineage using the following criteria: Erythroid (Ery): CD235a^+^CD45^-^CD11b^-^CD14^-^CD19^-^CD335^-^; Megakaryocytes (Mk): CD45^+^CD41^+^CD14^-^CD19^-^CD335^-^; Monocytes (Mono), CD45^+^CD41^-^CD14^+^CD11b^+^CD19^-^CD335^-^; B: CD45^+^CD41^-^CD14^-^CD19^+^CD335^-^; NK: CD45^+^CD41^-^CD14^-^CD19^-^CD335^+^; or not defined as any of these lineages (undefined). Additionally, a sample must contain ≥ 50 cells within the gate to be considered positive for a specific lineage. Cell counts were exported from Flowjo into R for quantification and visualization. Raw cell counts of B, NK, Mono, Ery and Mk cells were normalized to the total number of viable cells within each clone. Principal component analysis (PCA) was then conducted using the prcomp function with center = TRUE and scale. = TRUE. For log-scale visualization of output cell counts, zero counts were given a value of 0.5.

### Initial processing of single cell ATAC-seq data

The FL data and the adult BM dataset^28^ were run jointly through Cell Ranger ATAC version 1.2.0, using the human reference genome GRCh38 for alignment. The Cell Ranger output was processed using Signac v1.11^44^ and Seurat v4.4^45^. An initial Seurat object was created and divided to process the FL and adult BM cells separately.

FL cells were filtered to remove outliers for QC parameters (peak region fragments > 5000, peak region fragments < 55000, % reads in peaks > 40, ratio of reads from genomic blacklist regions < 0.005, nucleosome signal < 4 and TSS enrichment > 2.5), followed by term frequency-inverse document frequency (TF-IDF) normalization and singular value decomposition (SVD) for the top 25% of the features. The two FL samples were integrated using Harmony^25^. Harmony dimensions 2:20 were used for a first round of nearest neighbor analysis and unsupervised clustering (SLM algorithm, resolution = 0.7), and the clusters were used for per-cluster peak calling using MACS2^46^. The resulting peaks were subsetted to keep only chromosomes 1-22 to avoid artefacts due to different sexes of the donors, and peaks ≤ 20 or ≥ 10000 bp were removed. A new counts matrix was constructed using the MACS2 peaks and TF-IDF, SVD and Harmony were run again, using all features for SVD. Harmony dimensions 2:20 were used for clustering (resolution = 0.8) and Uniform Manifold Approximation and Projection (UMAP) generation (min.dist = 0.3, n.neighbors = 30, spread = 1).

The BM cells were processed in a similar manner, using the following QC parameter cutoffs: peak region fragments > 5000, peak region fragments < 60000, % reads in peaks > 45, ratio of reads from genomic blacklist regions < 0.005, nucleosome signal < 4 and TSS enrichment > 2.5. Additionally, cells not annotated in the original publication^28^ were excluded. For the final UMAP and clustering, resolution = 0.7 was used for clustering and min.dist = 0.2 for UMAP generation.

### Transcription factor motif analysis

Transcription factor (TF) motif activity was computed using chromVAR^26^ with the filtered (“version 2”) human TF motifs from the CIS-BP database^47^ available in the chromVARmotifs package. Differential motif activity between clusters was calculated using Seurat’s FindMarkers function (mean.fxn = rowMeans) and using average difference > 1.5 and adjusted p-value < 0.05 as cutoff values. For the dot plot in Figure 2, a maximum of 3 motifs per cluster were included, selected based on highest differential activity. Scores for sets of motifs were calculated using the AddModuleScore_UCell function from the UCell package^48^ on the chromVAR values, with maxRank = 200. When running PCA, the datasets were subsetted to contain clusters of interest and the top 150 motifs by variance were selected. The motif activities were scaled using Seurat’s ScaleData function and PCA was conducted using the prcomp function.

Transcription factor footprinting was investigated using Signac’s Footprint function with in.peaks = TRUE.

### Label transfer

To be able to perform the label transfer between adult BM and CS22 FL, a common peak set was created through merging overlapping peaks, removing peaks ≤ 20 or ≥ 10000 bp, and the new peaks were quantified in both datasets using the FeatureMatrix function. After TF-IDF normalization and SVD (top 95% of features), the FL dataset was projected onto the adult BM UMAP using Seurat’s FindTransferAnchors and MapQuery functions, using LSI dimensions 2:30.

### Differentially accessible peaks

Differentially accessible peaks were found using Seurat’s FindMarkers function with LR test and using the peak count per cell as latent variable. Peaks with absolute log2FC > 2 and p-adj < 0.05 were considered differentially accessible. To find motifs enriched in peaks differentially accessible between fetal and adult BM HSCs, we used Signac’s MatchRegionStats function to select a set of 20,000 background peaks among peaks that were accessible in at least 5% of HSCs from the two tissues compared. We then used the FindMotifs function and selected motifs with fold enrichment > 1.5 and p-adj < 0.05. Results were visualized using the ComplexHeatmap^49^ package with default clustering settings.

### Pseudotime analysis

The lymphoid trajectory for FL and adult BM combined was created through merging the two objects using a common peak set and combining the BM UMAP and the FL UMAP coordinates from the label transfer. Monocle3^50^ was then used to calculate pseudotime trajectories. For heatmap visualization of the activity of lymphoid-upregulated motifs along the trajectory, cells included in the trajectory were selected and the chromVAR values scaled across the subset (separately for FL and adult BM). The cell-motif matrix for adult BM was clustered using the hclust function and then ordered using dendsort (isReverse = T, type = ’min’). This row order was then used for both the adult BM and FL matrices to allow for easier comparison between the two.

When plotting chromVAR values or module scores as dots along pseudotime, a line was fitted to the individual cells using the geom_smooth function in ggplot2 with method = “gam” and formula = y ∼ s(x, bs = “cs”). ChromVAR values were scaled across the entire dataset prior to subsetting cells for plotting.

### Lineage affiliation

Lineage affiliation analysis was done using an R implementation of CellRadar (using data from HemaExplorer^51^ and also available at https://karlssong.github.io/cellradar/): For each gene set and cell type, the median expression value from the HemaExplorer data was extracted and the values were min-max scaled. Radar plots were created using the plotly package (https://plotly.com). Number of genes found in the HemaExplorer dataset/total number of genes for the different radar plots: Figure 4B: 53/64 for FL Ly-II, 53/57 for FL GMP-II, 61/82 for BM Ly-II, 75/78 in BM GMP-II; Figure S4B: 48/58 for FL Ly-I, 34/36 for FL GMP-I, 45/53 for BM Ly-I, 53/56 in BM GMP-I.

### Analysis of second trimester fetal liver

Publicly available single cell ATAC-seq data from LIN^-^CD34^+^CD38^-^ cells from three 18-21 pcw donors^5^ were downloaded. The raw data was processed using the nf-core atacseq pipeline v2.1.2^52^ using GRCh38 and a fragment file was generated using the sinto package (https://github.com/timoast/sinto). After an initial round of MACS2 peak calling (peaks filtered as described above for CS22 FL), the resulting Seurat object was filtered to keep cells that were included in the original publication for which 55000 > peak count > 5000 and TSS enrichment > 2.5, resulting in 3,515 cells. After normalization, Harmony integration, clustering and UMAP generation (dimensions 2:10, resolution = 0.5), peak calling was on a performed per-cluster basis, excluding clusters with too few cells, and the resulting peak set was used to generate a new integrated UMAP. The label transfer from adult BM to the 18-21 pcw dataset was performed as described above, except with LSI dimensions 2:10. The dataset was merged with the CS22 and adult BM datasets, as described above for the merge of CS22 and adult BM.

For the 12 pcw data (2 donors, a total of 3 replicates)^22^, the fragment files were downloaded and initial MACS2 peak calling was performed on each fragment file (for a total of 3 replicates) and replicates were filtered individually to remove low-quality cells. After this, processing was the same as for the 18-21 pcw data except that the top 25% of features were used for SVD, and Harmony dimensions 2:20 and resolution 0.6 were used for UMAP generation. To select progenitor populations comparable to the CD34^+^ CS22 and adult datasets from this CD45^+^ dataset, we first projected our CS22 FL dataset onto the 12 pcw UMAP (using Seurat’s FindTransferAnchors followed by MapQuery, using dims 2:20 of the LSI calculated on the top 25% of the merged peaks). We subsetted the 12 pcw dataset to keep only the clusters to which > 100 CS22 cells projected. We then projected this subset of 6,978 12 pcw cells onto the adult BM dataset and annotated them according to their predicted cluster identity.

### CITE-seq data analysis

The CS22 FL dataset from^21^ was merged with the adult BM dataset from^28^ and gene expression for genes and clusters of interest was visualized using Seurat’s DotPlot function. The cluster annotations from the two original publications were kept.

### Statistical analysis

For comparisons at the end points of pseudotime trajectories, the pseudotime was divided into 10 sections using the cut function in R. The 10^th^ section was selected to represent the final 10% of the trajectory in the FL, and for the lymphoid trajectory in the adult BM a range corresponding to the minimum and maximum values of the FL pseudotime range was selected. The cells in this range of the lymphoid trajectory were then subjected to statistical analysis. For module scores, (Ly-II score) - (GMP-II score) was calculated for each cell. The two sets of differences were then compared using the Wilcoxon rank sum test. For the TF motifs, the scaled (across the whole dataset) chromVAR value for CEBPA was subtracted from the scaled value of motifs of interest for each cell, and the distribution of differences was compared between FL and adult BM using the same test. The Bonferroni method was applied for p-value correction when performing multiple tests.

## Author contributions

S.P. and C.B conceptualized the study; S.P. and C.B performed the methodology; S.P. with input from K.N. and R.O. conducted the formal analysis; S.P., K.N. and M.S. conducted the investigation; V.T., G.K. and C.B. provided resources, S.P. and C.B wrote the original draft; C.B. with input from S.S. and G.K. supervised the study, C.B: acquired funds; S.P., V.T. and C.B. reviewed and edited the manuscript; all authors reviewed and approved the final manuscript.

## Acknowledgements

We thank the Lund Stem Cell Center FACS core facility at Lund University and the Center for Translational Genomics (CTG) at Lund University/SciLifeLab for technical assistance, Parashar Dhapola at Nygen Analytics AB for assistance with visualization using CellRadar, Agata Smialowska at SciLifeLab for valuable discussion, Ariana Calderón for assistance with illustrations, Roshanak Ghazanfari for assistance with *in vitro* culture experiments and David Bryder for providing valuable feedback on the manuscript.

The authors acknowledge the support from the Crafoord Foundation (20210777 C.B.) the Swedish Childhood Cancer Foundation (HFT2023-0006; PR2023-0065; C.B.), the Ragnar Söderberg Foundation (M34/18; C.B.), the Swedish Research Council (2023-02095 C.B. 2023-03188 G.K.), Swedish Cancer Society (23 3084 Pj C.B, 22 2475 Pj G.K.), Knut and Alice Wallenberg Foundation (2020.0210 G.K), and the national strategic research area grant StemTherapy.

## Supplementary Figure legends

**Figure S1. Developmental stage-specific lineage bias of hematopoietic stem cells (A)** Sorting strategy for single HSC (Lin^-^CD45^+^CD34^+^CD38^-^CD45RA^-^CD90^+^) *in vitro* culture experiments. Viable, single cells were gated on size and Lin^-^CD45^+^. Further gating is indicated in the figure. Red dots indicate the index sorted cells. FL (*left*), CB (*middle*) and adult BM (*right*). (**B**) Gating strategy to define lineage affiliation of *in vitro* colonies. Viable, CD45^+^ single cells were gated as indicated with arrows (*for definitions see method section*). **(C)** Number of colonies with output of B but no Mono, B and Mono, Mono but no B, or neither of these lineages, for all replicates combined. **(D-E)** PC analysis to visualize lineage representation at the single-clone level, (**D**) per developmental stage and (**E**) per lineage. The contours in D represent levels of estimated two-dimensional density. The color scales in E represent fraction of viable cells for the indicated lineage for each clone. (**F**) Number of colonies with output of one, two or ≥3 defined lineages, or no defined lineage, for all replicates combined.

**Figure S2. Quality parameters and cell state distributions for FL and adult BM ATAC-sequencing data (A)** Representative gating strategy, indicated with arrows, for the purification of Lin^-^CD45^+^CD34^+^ HSPCs from CS22 FL. **(B)** UMAP of CS22 FL cells after Harmony integration, colored by donor. (**C**) Peak counts per cell per Seurat cluster in CS22 FL. The median number of peaks for cluster #0 is shown as a line across all clusters. (**D**) The adult BM^28^ UMAP from Figure 2E colored by donor. **(E)** Peak counts per cell for each cell type in adult BM. The median number of peaks for the HSC cluster is shown as a line across all clusters. **(F)** *Left:* The FL cells projected onto the adult BM UMAP from Figure 2E and annotated by their predicted cell identities. *Right:* Prediction scores for the label transfer shown in Figure 1F-H. **(G)** Cell identity distribution for the FL and adult BM datasets as percent of total cells for each donor.

**Figure S3. Motif enrichment and occupancy in second trimester HSCs (A)** Second trimester (18-21pcw) HSPCs from^5^ projected onto the adult BM UMAP from Figure 2E and annotated by their predicted cell identities. **(B)** Prediction scores from the label transfer in (A), displayed per predicted cell identity. **(C)** Heatmap of TF motifs enriched in peaks differentially accessible between second trimester fetal BM and adult BM HSCs (fold change > 1.5; p-adj < 0.05). **(D)** Footprints of selected TFs from Figure 4C-D, for second trimester FL and fetal BM HSCs.

**Figure S4. Differentially accessible TF motifs and pseudotime trajectories for the FL and adult BM datasets (A)** Number of differentially accessible TF motifs for selected clusters compared to HSCs for CS22 FL and adult BM (mean z-score difference > 1.5; p-adj < 0.05). **(B)** Lineage affiliation of TF motifs with increased activity compared to the HSC cluster for the Ly-I and GMP-I clusters for CS22 FL (*left*) and adult BM (*right*). **(C)** Lymphoid pseudotime trajectory calculated on the combined FL and adult BM datasets, using the BM UMAP. *Left*: CS22 FL, *right*: adult BM. **(D)** Module scores for the TF motifs upregulated in adult BM Ly-II (*upper*) and GMP-II (*lower*) clusters, for select clusters in CS22 FL (*left*) and adult BM (*right*). **(E)** Difference between Ly-II cluster and GMP-II cluster scores per cell at the end of the fetal lymphoid trajectory, in CS22 FL and adult BM, used for the statistical calculation in Figure 4C (*see method section*). (**F**) Difference between PAX5 vs. CEBPA (l*eft*) and RUNX1 vs. CEBPA (*right*) TF motif activity per cell at the end of the fetal lymphoid trajectory, in CS22 FL and adult BM, used for the statistical calculation in Figure 4D (*see method section*). **(G)** Venn diagram of TF motifs up- or downregulated in FL or adult BM Ly-II cells compared to HSCs. (**H**) Second trimester (12pcw) FL cells from^22^, corresponding to HSPCs (*see method section*), projected onto the adult BM UMAP from Figure 2E and annotated by their predicted cell identities. **(I)** Prediction scores from the label transfer in (H), displayed per predicted cell identity.

