## Supplementary figures and images for "Chromatin Accessibility Shapes Developmental-Specific Lineage Plasticity in Hematopoiesis"

### Supplementary Figures 1-4

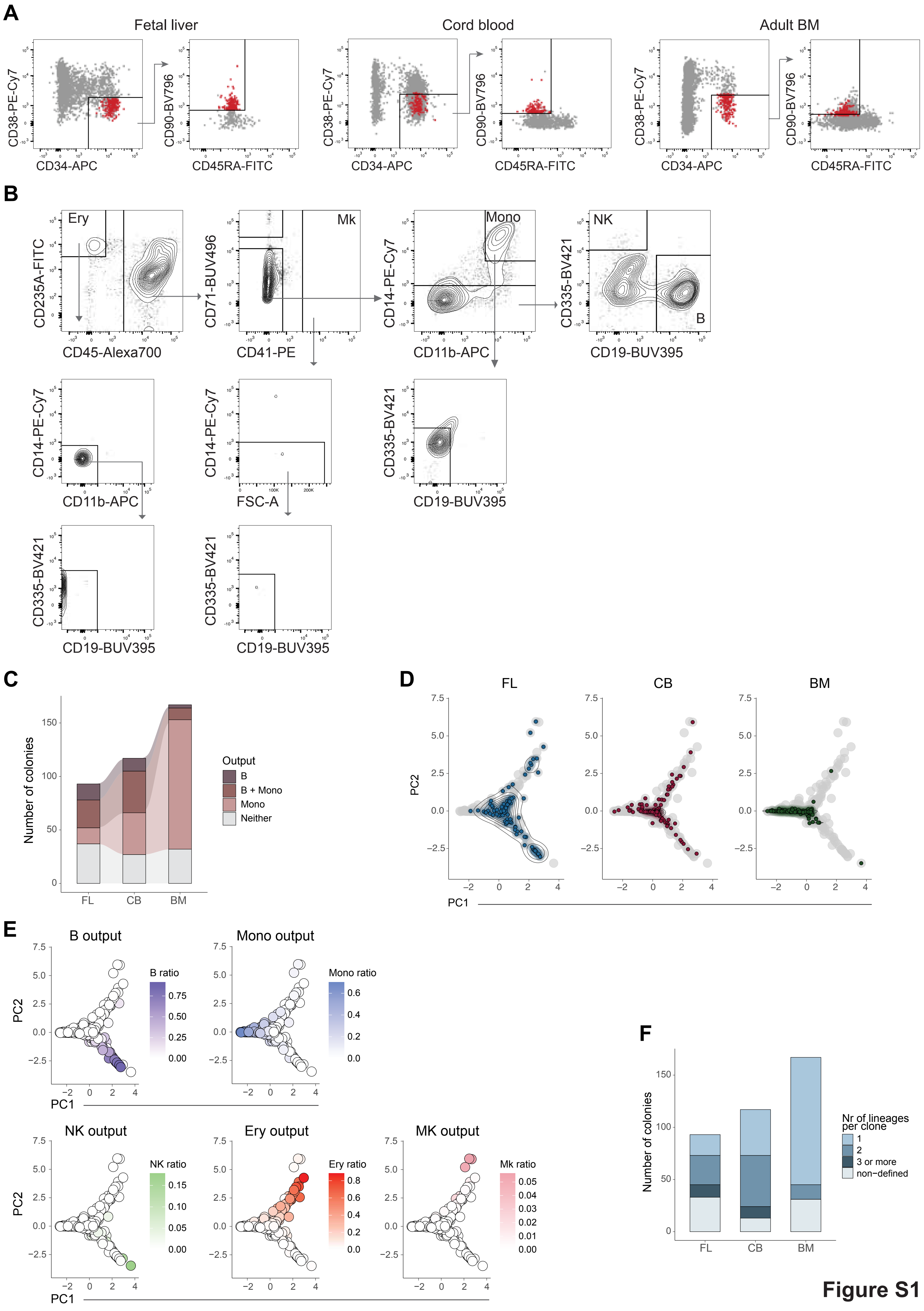

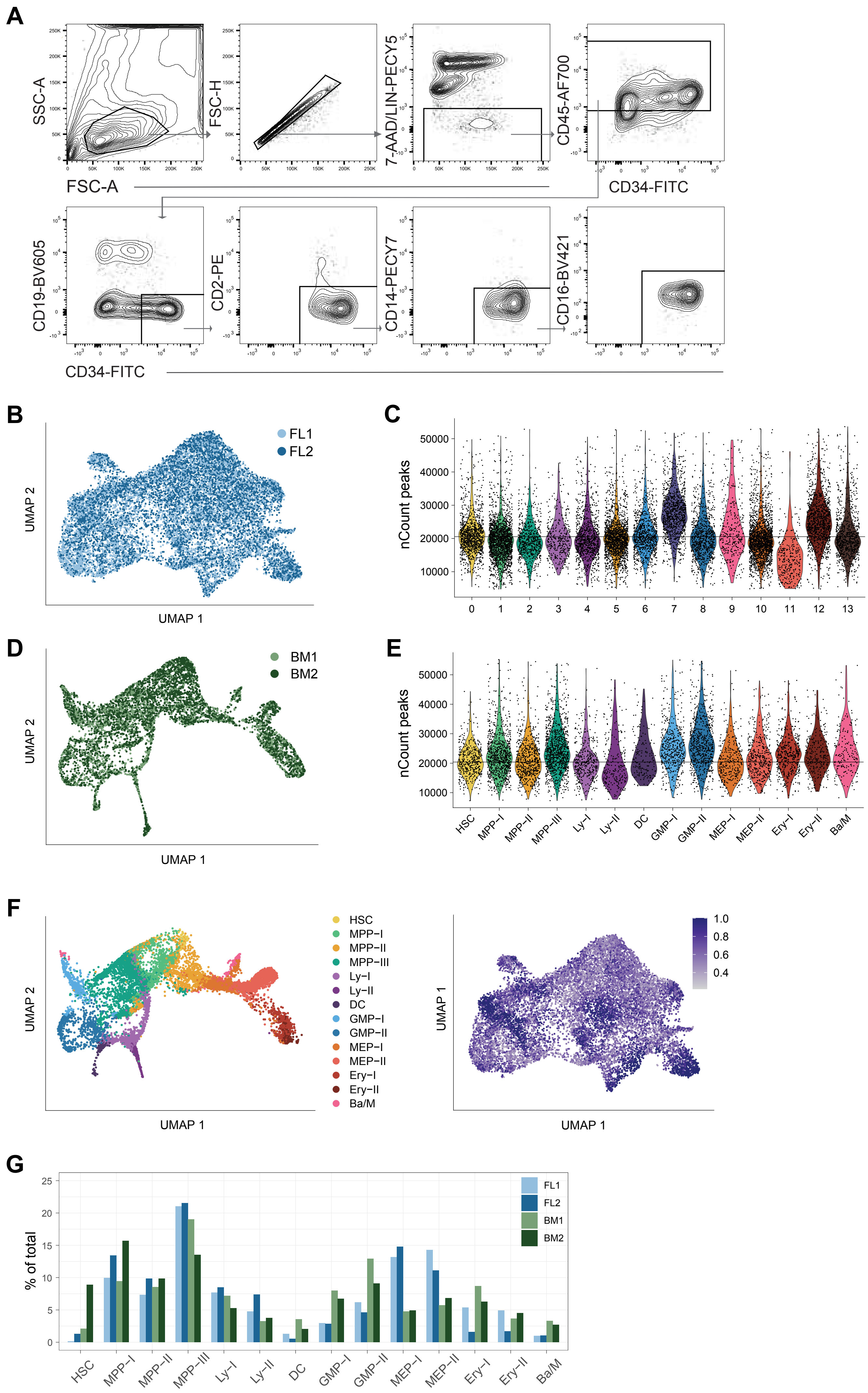

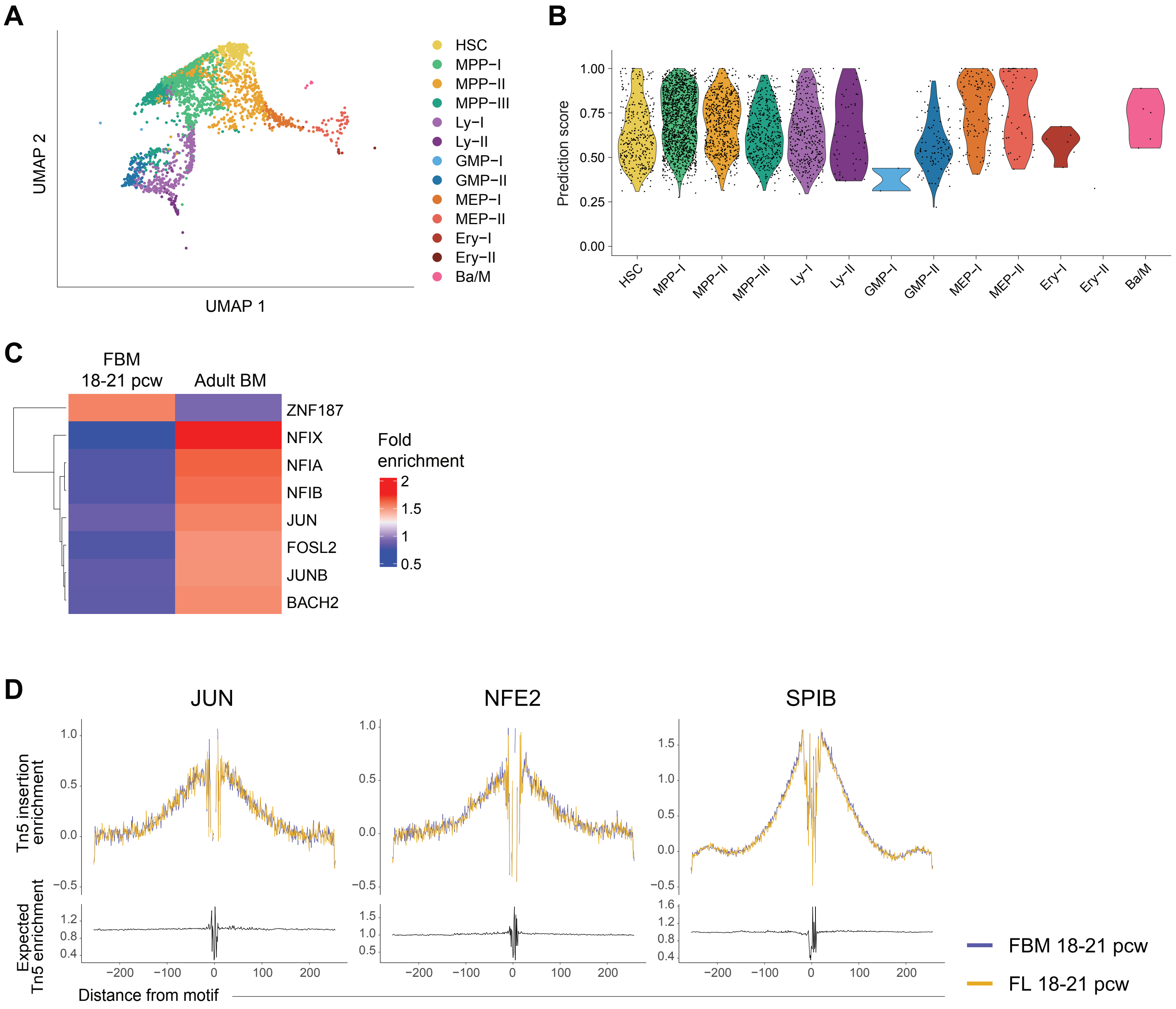

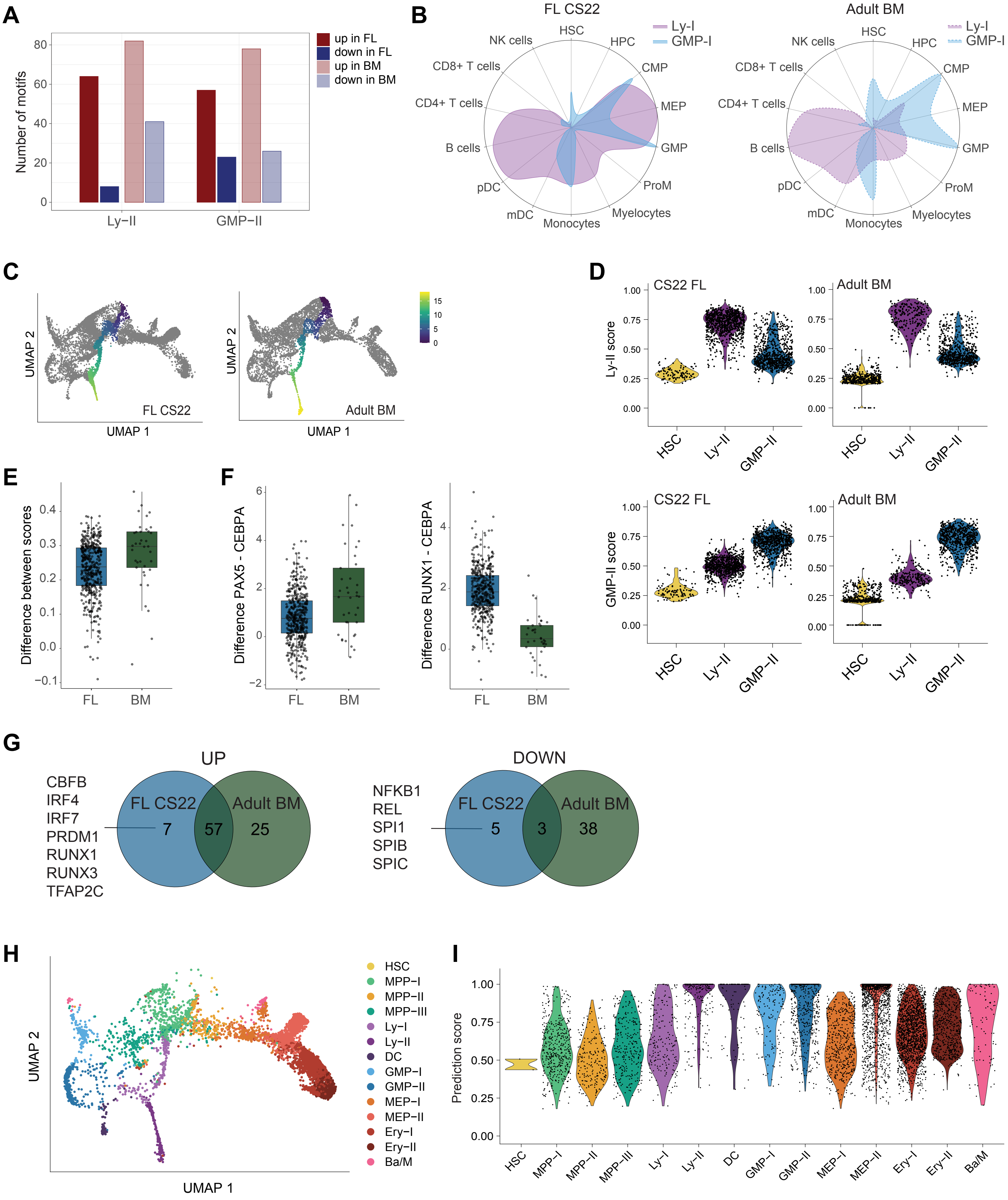
